# 5-aminolevulinic acid-photodynamic therapy ameliorates cutaneous granuloma by killing drug-resistant *Mycobacterium marinum*

**DOI:** 10.1101/2021.12.20.473605

**Authors:** Zhiya Yang, Yahui Feng, Zhiping Pang, Dongmei Li, Sisi Wang, Huiqi Chen, Mingze Jiang, Hongxia Yan, Tianhang Li, Hongjun Fu, Hubao Xiong, Dongmei Shi

## Abstract

**Background:** Although 5-aminolevulinic acid photodynamic therapy (ALA-PDT) has been extensively used to treat to various skin disease, the application of ALA-PDT on cutaneous infection caused by *Mycobacterium marinum* (*M. marinum*), especially by drug-resistant *M. marinum* is still not clear.

**Objectives:** We evaluated the therapeutic efficacy of ALA-PDT on *M. marinum in a* mouse infection model and tested its killing effect on *M. marinum in vitro*. Finally, we investigated the clinical effect of ALA-PDT in treating cutaneous granuloma caused by drug-resistant *M. marinum*.

**Materials and methods:** We isolated total 9 strains of *M. marinum* from patients and confirmed by morphological and molecular approaches. The strains were identified by anti-mycobacterial susceptibility. Then we evaluated the killing effect of ALA-PDT on *M. marinum in vitro* and in a mouse model to observe the antimycobacterial effect of ALA-PDT. Therapeutic efficacy was further assessed in patients with cutaneous granuloma caused by drug-resistant *M. marinum*.

**Results:** We demonstrated that the ALA-PDT directly killed *M. marinum in vitro*. The paws cutaneous lesions of mice caused by *M. marinum* were fully recovered 2 weeks after ALA-PDT treatment. However, there was no significant difference for immune cells in peripheral blood before and after ALA-PDT therapy. Finally, ALA-PDT proved to be effective in treat two patients with cutaneous infection caused by drug-resistant *M. marinum*.

**Conclusions:** The results suggest that ALA-PDT is effective in treating *M. marinum* cutaneous infections by killing *M. marinum* directly, independent of systemic immune responses. The data highlight the ALA-PDT as a promising therapeutic choice for *M. marinum* infection, especially for drug-resistant strains.

## Introduction

Cutaneous granuloma is a chronic skin inflammatory condition, which can be triggered by a wide spectrum of infectious agents including bacteria, viruses, fungi and other pathogens^[1]^. *Mycobacterium marinum* (*M. marinum*), belonging to non-tuberculous mycobacteria (NTM), is an important causative agent of infectious granulomas^[2]^. The disease is prevalent worldwide, with an incidence of 0.27/10^5^ in the United States and 0.04/10^5^ in France, as high as 6% on the island of Satowan in Cronesia^[3–5]^. But the incidence of *M. marinum* is still increasing rapidly and it has become a serious public health problem^[6–8]^.

Generally, *M. marinum* can cause human infection ranging from superficial to rarely disseminated or invasive infection even in immune-compromised patients^[9]^. The typical feature of superficial infection by *M. marinum* shows skin infectious granuloma^[10]^. Antibiotics is the primary choice for the treatment of patients with *M. marinum* skin infections. In accordance with the ATS/IDSA guidelines, currently ethambutol is recommended for the treatment of *M. marinum infection*, followed by rifampicin, clarithromycin and azithromycin^[11–13]^. Since *M. marinum* is a naturally drug-resistant organism, even a long course of antibiotics treatment is usually not effective ^[14]^. In a survey, 18% or 23% patients did not respond well to clarithromycin or a combination of ethambutol and rifampicin treatments, respectively, and relapse occurred in 24% of patients. Although the doxycycline or minocycline alone was reported to be effective in treating *M. marinum* infections ^[15–17]^, its side-effects and the need of long-term use make it difficult to treat patients clinically. Thus, it is urgently needed to find a new approach to kill *M. marinum* or enhance the antimicrobial activity of existing antibiotics^[18]^.

5-aminolevulinic acid photodynamic therapy (ALA-PDT) is a new, non-invasive therapeutic approach that combines the application of photosensitizers and light sources to damage selective targets through photodynamic reactions^[19]^. Topical drug application directly targets the lesion areas that reduces or minimizes the side effects on surrounding tissues^[20]^. The non-specific action of the released reactive oxygen species is less likely to induce any resistance mechanisms^[21, 22]^. In recent years, although it has been reported that ALA-PDT is effective in treating skin related diseases, its application in skin infections including *M. marinum* is not well studied ^[23–25]^.

In the present study, we evaluated the killing effect of ALA-PDT on *M. marinum in vitro* and a mouse model of skin infection. We demonstrated that ALA-PDT is effective against *M. marinum* infection, resulting in the improvement of skin lesions. In addition, *in vitro* experiments showed that ALA-PDT killed *M. marinum* directly. The mycobacterial burden in lesion area is significantly decreased after treatment with ALA-PDT. However, the systemic immune response did not alter significantly before and after treatments. Clinically, we treated 2 cases of *M. marinum* infection with ALA-PDT and the patients recovered fully within 1 month after treatment. The results suggest ALA-PDT is a novel approach in treating bacterial infections including *M. marinum* and highlight ALA-PDT as a promising therapeutic choice for future treatment of bacterial infections, especially drug-resistant pathogens.

## Materials and methods

### The isolation of *M. marinum* strains

The samples were collected from patients with cutaneous granulomas in Jining No. 1 People’s Hospital and inoculated on modified Middlebrook 7H10 agar media, chocolate plate and SDA at 30°C for 2 weeks. The modified Middlebrook 7H10 agar media were prepared as follows: agar was autoclaved at 121°C for 15 mins, the agar was incubated in a water bath at 56°C, and then added with the Oleic Albumin Dextrose Catalase Supplement (OADC) supplements.

### The identification of *M. marinum* by morphology and 16S rRNA genes sequencing

Colonies grown on Middle-brook 7H10 agar media were stained by fluorescent calcofluor white (CFW) as previously described^[26]^ and examined by fluorescence microscope. In addition, the isolates were also stained with acid-fast staining (the kit BA-4090B from Zhuhai Baso Biotechnology Company, China) as manufacturer instruction^[27]^ and then observed under light microscope.

For the identification of colonies, the genomic DNA was extracted by the guanidinium chloride method^[28]^. Briefly, the bacterial cells were treated by enzymatic digestion, mechanical disruption of the cell walls, and genomic DNA was extracted with phenol/chloroform/isoamyl alcohol 25:24:1. The DNA precipitates were then recovered by centrifugation and were dissolved in 1× TE buffer (10 mM TRIS, 1 mM EDTA) and were stored at −20°C before 16s sequencing. The 16s sequencing primers were described as follows^[29]^. RAC1 (5’ TCG ATG ATC ACC GAG AAC GTG TTC 3’) and RAC8 (5’ CAC TGG TGC CTC CCG TAG G 3’).

### ALA-PDT treatment of cutaneous *M. marinum* infection in a mouse model

First, we established a mouse model for cutaneous *M. marinum* infection as follows. *M. marinum* was diluted in 0.9% saline and each mouse paw was injected with 50μL (10^8^ CFU) *M. marinum* and infected for 2 weeks. Then, the mice were divided into 5 groups as follows: un-infected control group (A), infected only group (B), infected and treated with clarithromycin group (C) (gavaged with clarithromycin 2.6 mg per day for 2 weeks), infected and treated with rifampin group (gavaged with rifampicin 1.17 mg once time per day for 2 weeks) (D), and infected and treated with ALA-PDT group (E). In ALA-PDT group, each mouse was treated with ALA at a concentration of 20% for 3 hours followed by PDT irradiation at 100 J/cm^2^ using a 635-nm laser for 20 min once a week for 2 weeks.

### The analysis of systemic immune responses after ALA-PDT in mice *M. marinum* infection model

The blood was taken from the mice as described above and stored in an anticoagulant tube for the following experiments. Single-cell suspensions of peripheral blood were prepared, and immune cells were analyzed by flow cytometry with antibodies, including anti-CD3-PE, anti-CD4-FITC, anti-CD8-APC, anti-B220-FITC anti-NK1.1-PE, anti-CD11C-APC, anti-CD11b-FITC, anti-F4/80-PE and anti-CD86-CY7^[30]^. The cytokines levels in peripheral bloods including IL-12, IL-6, and IL-1β were assessed by ELISA.

### Antimycobacterial susceptibility test of the isolated *M. marinum* strains

Antimycobacterial susceptibility tests were performed by broth microdilution method as described ^[31]^. The bacterial suspension of each strain was prepared, and the turbidity of the suspension reached a density of 0.5 McFarland standard. An aliquot of 50 μL of stock suspension was added into 10 mL CAMHB (Thermo Fisher) with 5% OADC to achieve the final concentration (5×10^5^ CFU/mL), 100 μL of which were then transferred to each well of the 96-well plates for antibacterial susceptibility tests.

### The killing assay of ALA-PDT on *M. marinum*

The ALA solution was prepared by dissolving the ALA powder (Fudan Zhangjiang Company, Shanghai, China) in sterilized water to a final concentration of 20% (m/v) as stock solution. A series of working concentrations (20%, 10%, 5%, 2.5%, 1.25%, 0.625%, 0%) of ALA were prepared and 100 μL aliquots of each concentration was used as the photosensitizer. The bacteria were incubated with ALA in a 96-well plate for 30 min in the dark. Then, ALA was activated by the light source (630 nm LED, Omnibus, UK), and the output power was set at 100 J/cm^2^ for 20 min. The distance from the LED to the surface of the plates was 4 cm. The optimal ALA inhibitory concentration for each strain was determined by the absorbance at 600nm using a Multifunctional Microplate Reader (SynerghH1) at different time points.

### Patients with *M. marinum* cutaneous infections were treated ALA-PDT

Patients diagnosed clinically as skin granulomas caused by *M. marinum* were enrolled for the treatment. The lesions of patients were smeared with 20% ALA in dark for 3 hours followed by PDT irradiation at 100 J/cm^2^ using a 635-nm laser for 20 min. A total of 5 cycles of ALA-PDT treatment were applied for each patient at intervals of 10 days.

### Statistical analysis

One way ANOVA (and nonparametric or mixed) were used to analyze the effects of ALA-PDT Differences between or among groups in other experiments were assessed using the. Data were presented as means ± standard deviations. Values of P < .05 were considered statistically significant. GraphPad Prism 8.0 was used to generate figures.

### Ethics statement

This study was legal guardians of minors by Jining No. 1 People’s Hospital, Shandong, China legal guardians of minors. Written informed consent was obtained from all participants.

## Results

### The clinical isolates were identified as *M. marinum* by morphological and molecular approaches

To identify *M. marinum*, we used chocolate blood plate to culture the bacteria at 30°C. When the plate was kept in the dark, the colony of *M. marinum* was usually smooth and white but became yellow upon exposure to light (photochromogenic). The colonies of *M. marinum* is neat or slightly irregular (Figure 1A and 1B). We isolated a total of 9 strains suspected *M. marinum* from the local hospital and all the strains were characterized as unmoved polymorphous rod (1.0 to 4.0μm by 0.2 to 0.6μm) with true branches, which are undistinguished from other *Mycobacterium* spp. (Figure 1C). The 9 strains were all positive for CFW and acid-fast staining under fluorescence and light microscopy respectively (Figure 1D). In addition, 16S rRNA sequencing analysis confirmed all these isolates are *M. marinum* (Table 1 and Figure 1E). Through anti-bacteria susceptibility test, we identified 2 among 9 strains were multi-drug resistant *M. marinum* strains.

**Figure 1.**
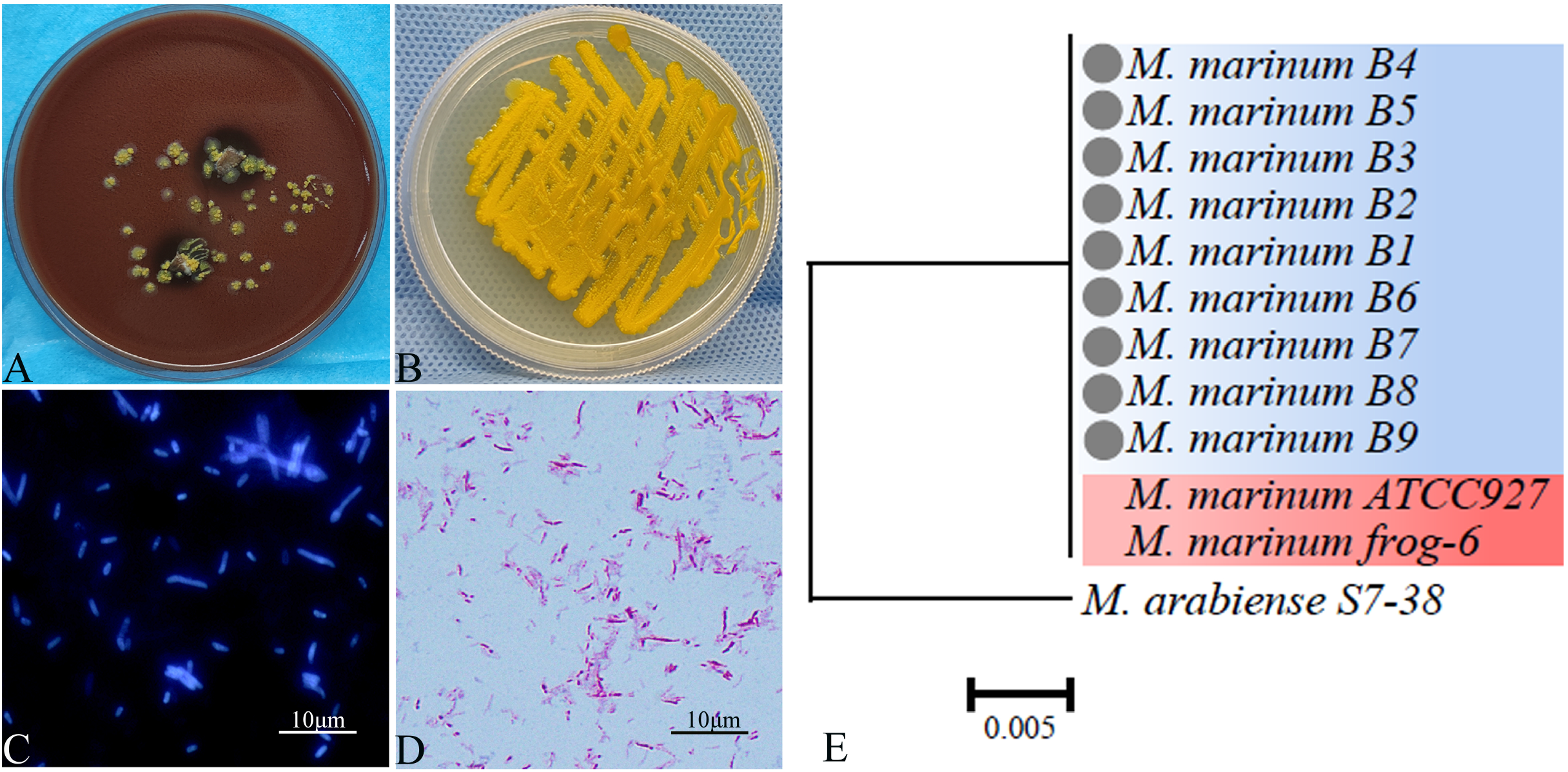
Identification of *M. marinum* isolated from patients’ lesions. A, the samples from patients’ lesions were inoculated and cultured on chocolate medium for 2 weeks, then dark yellow colonies were seen as exposed to light. B, the colony grew on chocolate medium was transferred to modified Middle-brook 7H10 agar for 2 weeks. C, *M. marinum* was stained with calcium fluorescent white and observed under fluorescence microscope. D, *M. marinum* was stained with acid-fast staining and observed under light microscope. E, a phylogenetic tree was constructed using the maximum likelihood method based on the 16S sequencing. *M. marinum* ATCC927 severed as standard strain and *M. arabiense* was as an outgroup strain.

**Table 1.** Clinical isolates of *M. marinum* from skin infected patients

| NO. | Age | Infected lesion | Molecular identification | GenBank NO. | Identify |
| --- | --- | --- | --- | --- | --- |
| 1 | 29 | Left hand | *Mycobacterium marinum* | MZ540348 | 100% |
| 2 | 38 | Right arm | *Mycobacterium marinum* | MZ540350 | 100% |
| 3 | 21 | Right leg | *Mycobacterium marinum* | MZ540351 | 100% |
| 4 | 24 | Left arm | *Mycobacterium marinum* | MZ540352 | 100% |
| 5 | 63 | Right arm | *Mycobacterium marinum* | MZ540356 | 100% |
| 6 | 53 | Left arm | *Mycobacterium marinum* | MZ540357 | 99.71% |
| 7 | 38 | Left arm | *Mycobacterium marinum* | MZ540359 | 99.93% |
| 8 | 50 | Right arm | *Mycobacterium marinum* | MZ540361 | 100% |
| 9 | 48 | Left arm | *Mycobacterium marinum* | MZ540362 | 100% |

### The surprising effectiveness of ALA-PDT in the treatment of lesions infected by *M. marinum* in a mouse model

At first, we established a mouse model of skin infection with *M. marinum*. The paws of mice infected with *M. marinum* (10^8^ CFU bacteria per mouse) and the paw of mice showed redness, swelling, rupture and pus flow 2 weeks after infection (Figure 2). Then, the mice were divided into 5 groups and were treated as follows: untreated group (A), infected only group (B), oral clarithromycin treated group (C), oral rifampicin treated group (D), and ALA-PDT treatment group (E).

**Figure 2.**
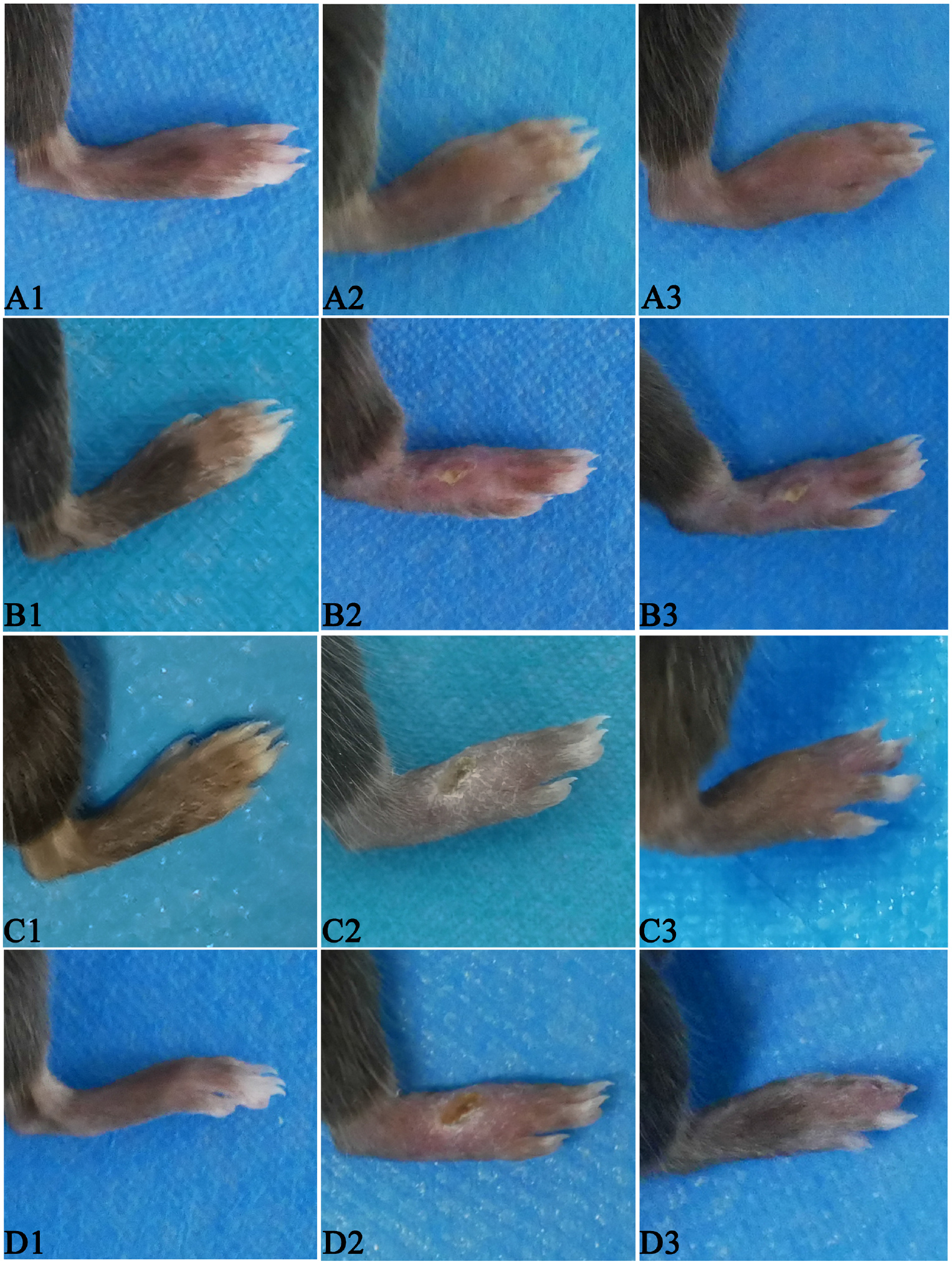
ALA-PDT improved mouse paw lesions caused by *M. marinum* infection. Mice were infected with 10^8^ CFU *M. marinum* for 2 weeks. Mice were infected and treated with PBS (A1-A3), or treated with rifampicin (B1-B3), or treated with clarithromycin (C1-C3), or treated with ALA-PDT (D1-D3) for 4 weeks. Images of mouse paws before and after treatment.

After 2 weeks of treatment, the results showed that clarithromycin and rifampicin were not effective in treating *M. marinum* infection, instead developed severe side effects result in weight loss. Surprisingly, in ALA-PDT treatment group we found that crusts appeared on the paws after one week of treatment, and the redness and swelling subsided significantly after two weeks of treatment (Figure 2). The mice recovered fully after four weeks of treatment, as confirmed by bacterial loads (Figure 2). The results suggest that ALA-PDT is effective in treatment of *M. marinum* caused skin infection.

### The effectiveness of ALA-PDT in treating *M. marinum* infection was independent of immune response

To investigate if immune response is involved in the treatment of *M. marinum* infection by ALA-PDT, we analyzed immune cells before and after treatment. The results showed that there were no significant changes for T cells, B cells, NK cells and monocytes (Figure 3). The level of cytokines in peripheral blood including IL-12, IL-6, and IL-1β were not detectable even after paws skin infection (data not shown). The results suggest that ALA-PDT does not activate systemic immune responses and its effectiveness in the treatment of *M. marinum* infection is independent of immune responses.

**Figure 3.**
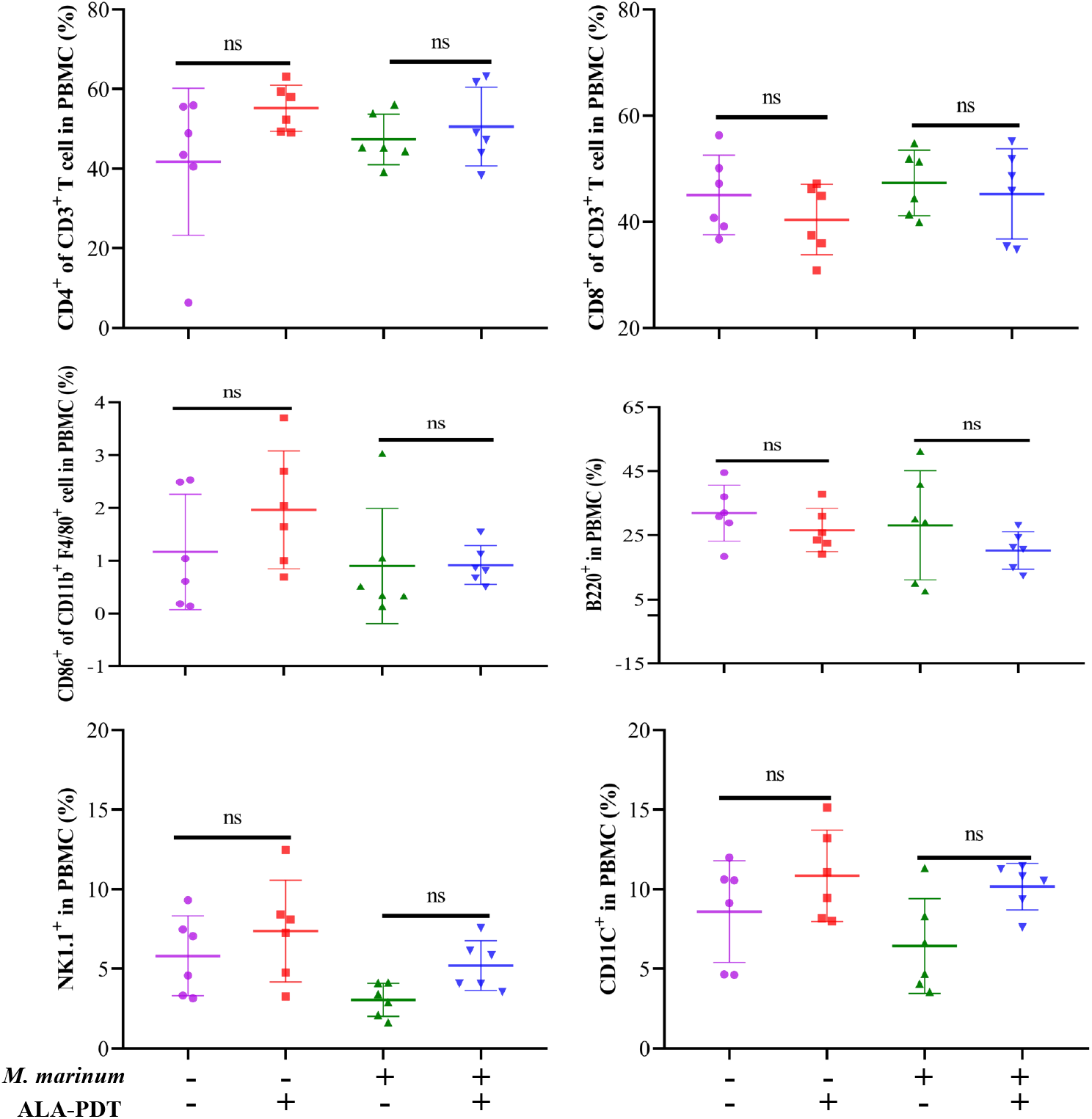
There were no significant changes for immune cells before and after ALA-PDT treatment. Mice were infected with 10^8^ CFU *M. marinum* for 2 weeks and then treated with ALA-PDT for 4 weeks. Mice were then sacrificed, and blood cells were collected and analyzed by flow cytometry. The results were representative of 3 similar experiments.

### ALA significantly inhibited drug-resistant mycobacterial growth in culture medium

To test the effect of ALA-PDT on drug-resistant *M. marinum* strains, we diluted ALA to various concentrations to determine the optimal working concentration in suppressing drug-resistant mycobacteria. The results showed that mycobacterial growth was significantly inhibited by ALA-PDT treatment when compared with *M. marinum* growth only. Interestingly, ALA at concentration above 1.25% ALA-PDT showed significant inhibition of *M. marinum* growth, while ALA concentration below 0.625% lost its ability to inhibit *M. marinum* growth (Figure 4A and 4B). In addition, to confirm the above results, the culture mediums containing various ALA concentration were further cultured on Middlebrook 7H10 (M7H10) agar media for mycobacteria growth. The results showed that there were yellow colonies at ALA concentration below or equal 0.625%, however there was no colony of *M. marinum* with ALA concentration above or equal to 1.25% (Figure 4C). Furthermore, ALA alone or PDT alone had no effect in suppressing *M. marinum* growth. Then, we used 5% ALA concentration to test remaining 7 strains (not drug-resistant), and showed similar results. The results suggest ALA-PDT effectively kills *M. marinum*.

**Figure 4.**
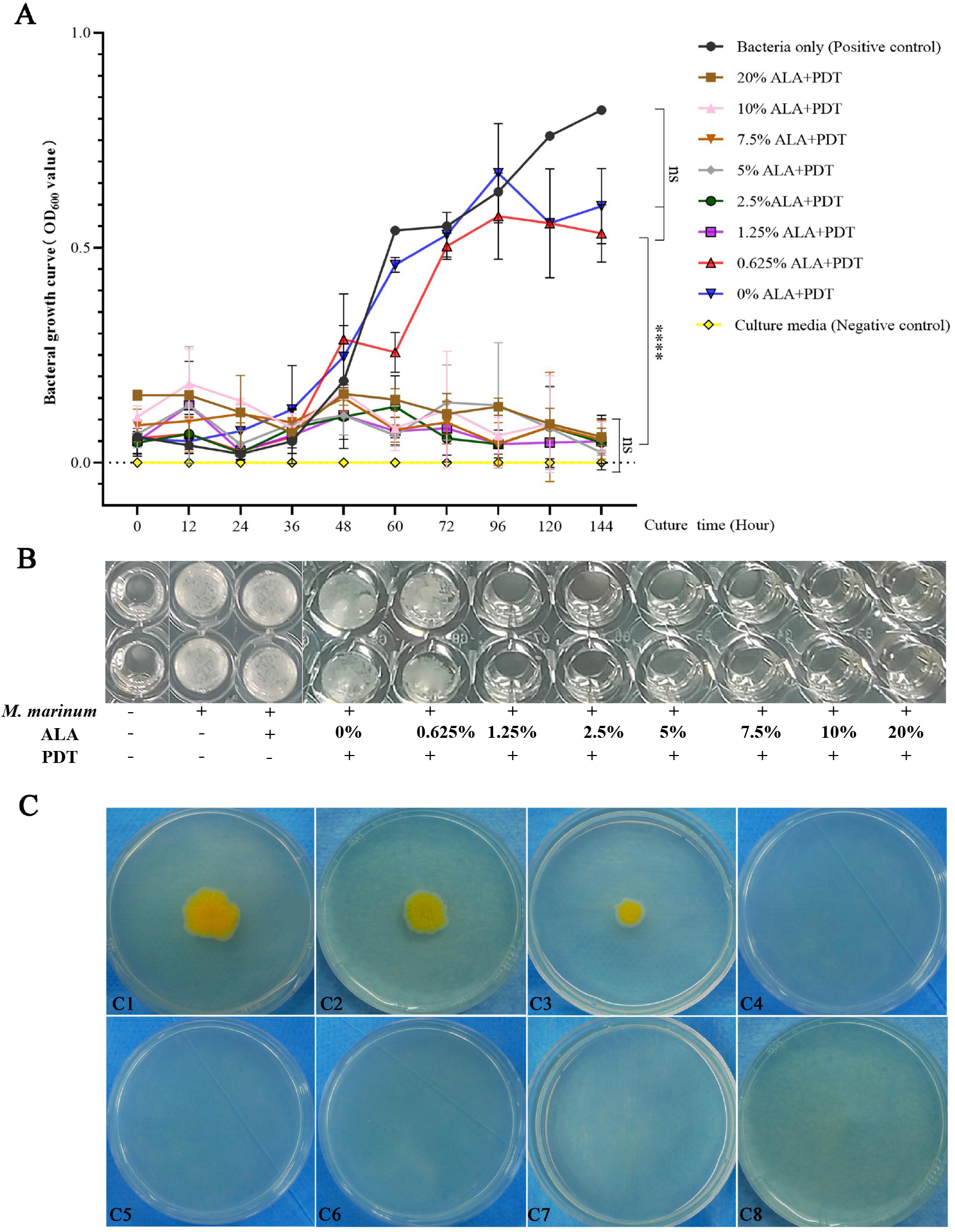
The effects of ALA-PDT on *M. marinum*. The bacterial growth curve for 144 hours. *M. marinum* was suspended into CAMHB medium at concentration of 5×10^5^ CFU/ml, added with ALA at different concentrations (20%, 10%, 5%, 2.5%, 1.25% and 0%), and the bacteria were cultured for 30 min in dark followed by PDT radiation for 20 min. A, the bacterial growth was measured at OD_600_ and the curve was illustrated. B, the images of turbidity of the above wells. C, the aliquots from the above wells were inoculated on modified Middle-brook 7H10 and cultured for 2 weeks. The colony formation was monitored.

### The patients with lesions caused by drug-resistant *M. marinum* fully recovered by ALA-PDT treatment

The above results demonstrated that ALA-PDT can directly kill *M. marinum* **a**nd improve lesions in mouse *M. marinum* infection model. Next, we wanted to treat patients with cutaneous lesions caused by drug-resistant *M. marinum*. We selected 2 patients that were initially suspected of drug-resistant *M. marinum infection*. The patients were given routine physical examinations, along with routine blood work. Urine and fecal examination results were within normal ranges. Hepatitis B, hepatitis C, syphilis, and HIV were negative, and Chest X-rays were normal. The patients underwent histological examination, and the biopsied specimens were characterized as infectious granuloma (Figure 5A and Figure 5D). In addition, the histological specimens were negative for both PAS and acid-fast staining (Figure (5B, 5C,5E and 5F)). Based on the clinical symptom and laboratory tests, the patients were diagnosed as cutaneous granuloma caused by drug-resistant *M. marinum* and were treated with ALA-PDT for 5 times at interval of 10 days.

**Figure 5.**
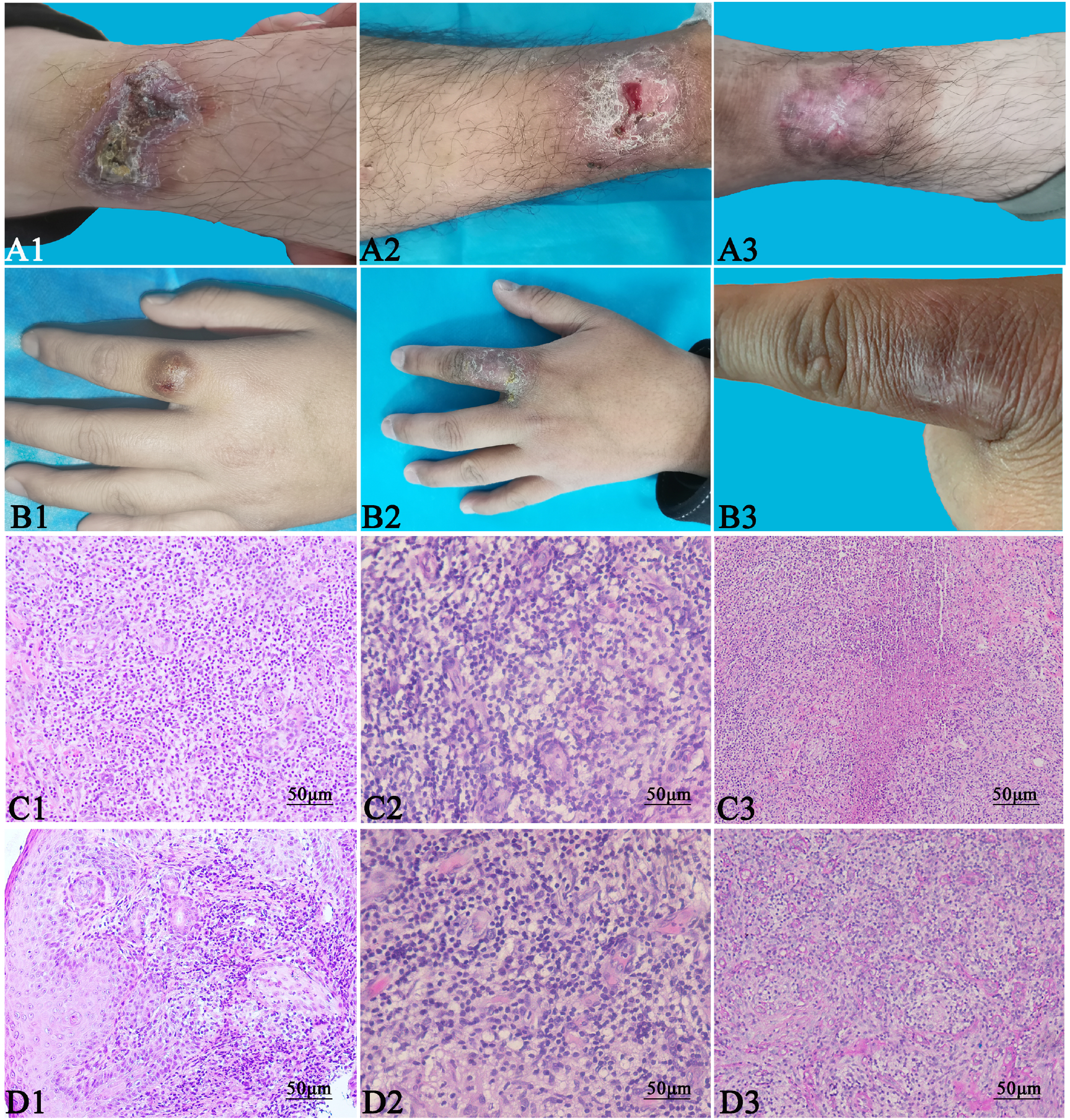
The patients with infectious granuloma by drug-resistant *M. marinum* were treated with ALA-PDT. The patients with infectious granuloma by drug-resistant *M. marinum* were enrolled for treatment by ALA-PDT for 5 times at interval 10 days. Clinical recovery of lesions caused by drug-resistant *M. marinum*. A1-A3 represent the treatment stage of patient No.1: before treatment (A1), after treatment for 5 times (A2), or 2 months follow-up after treatment (A3). B1-B3 represent the treatment stage of patient No. 2: before treatment (B1), after treatment for 5 times (B2), or 2 months follow-up after treatment (B3). C1-C3 represent the histological examination of patient No.1: HE staining (C1, original magnificent 200×), acid-fast staining (C2, original magnificent 200×), or PAS staining (C3, original magnificent 200×) for *M. marinum*. D’-D3 represent the histological examination of patient No. 2: HE staining (D’, original magnificent 200×), acid-fast staining (D2, original magnificent 200×) or PAS staining (D3, original magnificent 200×) for *M. marinum*.

The patients responded to ALA-PDT, showing that the lesions disappeared within 7 weeks. The patients had no recurrence at the 2-month follow-up (Figure 5G–Figure 5L). The results suggest that ALA-PDT is clinically effective for cutaneous granuloma caused by drug-resistant *M. marinum*, highlighting the importance of ALA-PDT as a primary treatment choice for skin infection caused by drug-resistant *M. marinum*.

## Discussion

*M. marinum* is ubiquitously waterborne opportunistic pathogen, which can also be found in other different environments^[32, 33]^. In the recent decades, the incidence of *M. marinum* infection has been increasing steadily^[34–36]^. The human infection cases were most commonly reported in coastal areas, but the patients we admitted to our hospital were from the inland area. Interestingly, all the patients had a history of trauma or long-term exposure to water or fish transported from the seaside.

Although the infection of *M. marinum* is mostly confined to the skin, systemic dissemination of infection is rare and has only been reported in immune-compromised patients and the diagnosis is difficult due to the lack of specific symptoms^[37–39]^. We found that *M. marinum* did not grow at 37°C but grew well at 30°C on chocolate medium, which could be reason why *M. marinum* only cause limited infection to the skin but not disseminated infection.

Currently, the clinical therapy for treating *M. marinum* cutaneous infection include antibiotic and surgery^[40]^. However, *M. marinum* has a natural multidrug resistance property which may be due to the long-term use of antimycobacterial drugs in clinical practice^[41]^. The appearance of drug-resistant strains of *M. marinum* strains make the treatment become more difficult. Thus, the new therapeutic strategies are required to improve clinically treatment effect and promote the patient’s recovery.

5-aminolevulinic acid photodynamic therapy (ALA-PDT) has been extensively used for the treatment of premalignant lesions or solid tumors^[42, 43]^. In recent years, ALA-PDT has been expended to treat various skin diseases, including skin infections caused by bacteria, viruses, fungi and other microorganisms^[23]^. However, its application in skin infection caused by *M. marinum* is not clear. In the present study, we used ALA-PDT to treat *M. marinum* infection in a mouse model and treated 2 patients with drug-resistant *M. marinum* infections, the results showed that ALA-PDT is effective in killing mycobacteria resulting in the full recovery of infected mice as well as clinical patients. Although, we have shown ALA-PDT is effective in treating drug-resistant *M. marinum* infections, it is still not clear whether ALA-PDT and antibiotics have synergistic effect in controlling *M. marinum* infections. We will use ALA-PDT together with antibiotics to address this question in our future study.

Although ALA-PDT was initially shown to have a therapeutic effect in skin infectious diseases, no detailed studies have been conducted to investigate its mechanisms. In the present study, we clearly demonstrated that ALA-PDT can kill *M. marinum* directly, independent of systemic immune response since we did not find any changes of immune cells before and after ALA-PDT treatment in mouse model. However, the detailed mechanism for ALA-PDT to kill *M. marinum* is not defined yet. It is generally believed that ALA-PDT can kill bacteria by effectively disrupting the membrane and wall of bacteria, resulting in bacterial death by leakage of content^[44]^. Bertoloni G el al. reported that ALA-PDT can damage both single and double-stranded DNA of bacteria in Gram-positive or negative bacteria after ALA-PDT in experimental settings^[45]^. Bartolomeu M et al. suggested that ALA-PDT causes inactivation of virulence factors through its effect on bacterial virulence factors^[46]^. Our future studies will focus on research concerning the molecular mechanisms for ALA-PDT to kill *M. marinum*.

In summary, we demonstrated that the ALA-PDT directly killed *M. marinum in vitro* and *in vivo*. The cutaneous lesions caused by *M. marinum* on mice paws were fully recovered after ALA-PDT treatment independent of immune responses. In addition, we clinically used ALA-PDT to treat drug-resistant *M. marinum* skin infection and the results showed that ALA-PDT is clinically effective for cutaneous granuloma caused by drug-resistant *M. marinum*, suggesting the importance of ALA-PDT as a primary treatment choice for skin infection caused by drug-resistant *M. marinum*.

## Acknowledgments

This work was supported in part by grants from the National Natural Science Foundation of China (NM 81773337), China Medical Fungi Alliance, (CMFA-2021-06) and ‘Qihang’ Plan of Jining No. 1 People’s Hospital (2021-QHM-026).

## Author Contributions

Zhiya Yang and Yahui Feng contributed equally to this work. Author order was determined by drawing straws. All authors contributed extensively to the work presented in the manuscript. DM S and H X conceived and designed the study. ZY Y, YH F, SS W, ZP P, HQ C and MZ J performed the experiments; DM L and ZY Y analyzed the data; TH L, HY Y and HJ F provide clinical cases DM S and ZY Y wrote and revised the manuscript.

